# Structures of Multiple Peptide Resistance Factor from *Pseudomonas aeruginosa*

**DOI:** 10.1101/2024.08.21.608925

**Authors:** Shaileshanand Jha, Kutti R. Vinothkumar

## Abstract

The aminoacylation of lipid head group in many bacteria is carried out by bi-functional enzymes called MprF, which encode for a soluble synthase domain that typically transfers lysine or alanine from a tRNA to lipid head groups, and the modified lipid is translocated across the leaflets by a transmembrane domain. This modification of the lipids probably evolved to adapt to the environment where the microbes reside. Here, we describe the cryoEM structures of MprF enzyme from *Pseudomonas aeruginosa* revealing a dimeric enzyme with a distinct architecture when compared with the homologous Rhizobium enzymes and validate this arrangement with biochemical analysis. The cryoEM maps and the models in detergent micelle and nanodisc reveal a conformational change of the terminal helix of the synthase domain, highlighting the dynamic elements in the enzyme that might facilitate catalysis. Several lipid-like densities are observed in the cryoEM maps, which might indicate the path taken by the lipids and the coupling function of the two functional domains. Thus, the structure of a well-characterised PaMprF lays a platform for understanding the mechanism of amino acid transfer to a lipid head group and subsequent flipping across the leaflet that changes the property of the membrane.

## Introduction

The role of transfer RNA (tRNA) as an adaptor molecule in translation is well studied and beyond this more common role, charged tRNAs also fulfil other roles such as donating amino acids to lipids or carbohydrates and, in bacteria these form important processes in cell wall biosynthesis and antibiotic resistance (Katz et al, 2016; RajBhandary & Söll, 2008). One such role for tRNA is found in the aminoacylation of lipid head groups, which not only allow microbes to adapt to the environment but also to counter the activity of small cationic peptides produced by a number of organisms (Fields & Roy, 2018; Ernst & Peschel, 2019; Slavetinsky *et al*, 2017). Organisms in particular microbes, to survive and grow in a competitive environment produce a number of small molecule metabolites that act on various cellular processes. Some of these molecules are the small peptides, which use the common property of the charge on the cellular membrane to kill other cells. These are collectively called cationic antimicrobial peptides (CAMPs), which are released by the host including eukaryotes and bind to the membranes and among other things promote pore formation (Scott et al, 1999; Michael Zasloff, 2002).

Naturally, microbes develop resistance to such molecules and in several pathogenic microbes as well as some archaea, one of the resistance mechanisms to CAMPs is mediated by a bifunctional enzyme called Multiple peptide resistance factor (MprF). The family of MprF enzymes typically have two domains: a cytosolic synthase domain that allows binding of a charged tRNA and transfer of amino acid, typically a lysine or alanine (or arginine), to the head group of a lipid on the inner leaflet (Peschel et al, 2001). The transmembrane (TM) domain performs the flipping of the lipid to the outer leaflet to introduce modified lipid on the membrane surface, thereby rendering the CAMPs to be ineffective (Ernst et al, 2009; Fields & Roy, 2018).

The lipid molecule that is commonly transferred with an amino acid is phosphatidyl glycerol (PG) but cardiolipin (CL) can also act as a substrate. MprF are diverse enzymes both in their substrate specificity and the product generated. The enzyme from *Corynebacterium glutamicum*, which synthesises alanyl diacylglycerol using AlaDAGS and, LysX from *C. pseudotuberculosis*, which is responsible for the synthesis of lysine PG and lysine diacyl glycerol are extremely specific (Gill et al, 2023). While, LysPGS from *Lysteria monocytogenes* transfers lysine to PG as well as to CL (Thedieck et al, 2006). MprF homologs from few other bacteria are promiscuous in transferring different amino acids to lipid molecules. These include MprF from *Bacillus subtilis*, which uses ala – tRNA as well as lys - tRNA as substrates for aminoacylation, and MprF2 from *Enterococcus faecium* that transfers arginine to PG in addition to alanine and lysine (Roy & Ibba, 2009). In contrast, *Clostridium perfringens* has two separate enzymes for synthesising AlaPG and LysPG (Roy & Ibba, 2008). When these MprFs are expressed in heterologous expression systems such as *E. coli* (where this enzyme is absent), modified lipids are synthesised and have been shown to decrease the effect of CAMPs, to provide resistance against antibiotics such as daptomycin, β-lactams, as well as aminoglycosides (Arendt et al, 2012; Nishi et al, 2004). In addition, there are reports of several gain of function mutations that have been acquired in antibiotic resistant strains of various pathogenic microbes including Methicillin resistant *Staphylococcus aureus* (MRSA) (Ernst & Peschel, 2019). Thus, characterisation of these enzymes is interesting in many aspects, including the nature of recognition of the specific tRNA by the synthase domain, mechanism of transfer of amino acid to the lipid head group as well as lipid transfer from one leaflet to the other.

Structurally, the soluble synthase domains from *Pseudomonas aeruginosa* and *Bacillus licheniformis* were the first to be characterised a decade ago revealing two GCN5-related N-acetyltransferase (GNAT) fold in tandem. Additionally, the Bacillus enzyme was solved with substrate analogue L-lysine-amide identifying a potential substrate binding site (Hebecker et al, 2015). The structure of a close homologue FemX from *Weissella viridescens,* another ala-tRNA transferase involved in peptidoglycan synthesis, revealed the peptidyl RNA conjugate binding site (Fonvielle et al, 2013). Together, these studies have been used to propose the possible binding sites for the aminoacylated acceptor stem arm of tRNA and a possible cavity for binding of the PG molecule. In 2021, the full-length MprF structure of *Rhizobium tropici* by cryoEM as well as the synthase domain by X-ray crystallography was determined (Song et al, 2021) and another cryoEM structure from *Rhizobium etli* has been deposited (EMD-31445). These structures provide information on the relative position of the membrane and the soluble domain, the buried lipid molecules in the flippase domain and the possible mechanism of amino acid transfer and lipid flipping from the inner to the outer leaflet. Perhaps, the interesting features of the full-length structures from *Rhizobium sp.* are the arch like assembly of the monomers, the dimeric interface and the flexibility of the soluble domain. The diversity in these enzymes for their substrates and products and their importance in resistance to peptide-like drugs prompts further characterisation of homologs from other organisms to understand their architecture and substrate specificity.

Here, we describe cryoEM structures of MprF from *Pseudomonas aeruginosa*, a well characterised enzyme that synthesises and flips AlaPG (Klein et al, 2009), which lacks a positive charge but confers resistance to CAMPs as well as various other anti-microbial compounds (Slavetinsky et al, 2012; Roy & Ibba, 2008; Arendt et al, 2012). The structures of PaMprF determined both in detergent micelle as well in nanodisc reveals a more compact dimer and a distinct dimer interface with respect to the homologous Rhizobium structures. Biochemically, we show that the dimer interface observed in PaMprF is preserved in the membrane and the choice of the detergents affects the stability of the enzyme. Few lipid-like densities are observed in the core of the flippase domain, along with a number of other partially or fully ordered lipid molecules in the membrane domain providing an explanation for the path taken by the lipids before or after modification to be assimilated with bulk lipids.

## Results

### CryoEM maps of PaMprF in detergent micelle and nanodisc

The polypeptide of PaMprF was expressed and purified as a fusion protein with a cysteine protease domain (CPD) and a His-tag (Shen et al, 2009). The tag was cleaved by inducing the CPD activity with inositol-6-phosphate (IP_6_). Extraction of the enzyme from the membrane with lauryl maltose neopentyl glycol (LMNG) and subsequent exchange to glyco-diosgenin (GDN) provided the best profile in size-exclusion chromatography. Peak fractions were pooled, concentrated and this sample was used for cryoEM studies. Reference-free 2D classification of PaMprF in GDN revealed a dimeric enzyme with different views and a general flexibility of the soluble synthase domain (Figure S1 A and B). Multiple rounds of 3D classification in Relion and heterogenous refinement in CryoSparc (Kimanius et al, 2021; Punjani et al, 2017; Scheres, 2012) were used to enrich the population of particles that had well resolved transmembrane domain (TMD). This revealed two major populations, one where both the soluble synthase domains were observed and the other where only one of the soluble domains was ordered (Fig S1C). The particle from these populations were refined and reconstructed without the application of symmetry or with C2 symmetry for the particles with both the soluble domains ordered. The overall resolution of the maps is between 3.3-3.4 Å (Figure 1A and Figure S1D, Table 1). As expected, the local resolution of the map reveals higher resolution at the core of the dimer, the peripheral TM helices at ∼4 Å and the soluble domain between 3.5-5 Å with well-resolved regions at the membrane interface (Fig S1E and S4A). The reconstructions of the PaMprF with ordered soluble domains in both monomers using C1 or C2 resulted in similar resolutions (3.4 Å and 3.3 Å respectively) with no major difference in the structure and we have used the C2 symmetrized map for the large part of the analysis (Figure S1 and Table 1).

**Figure 1:**
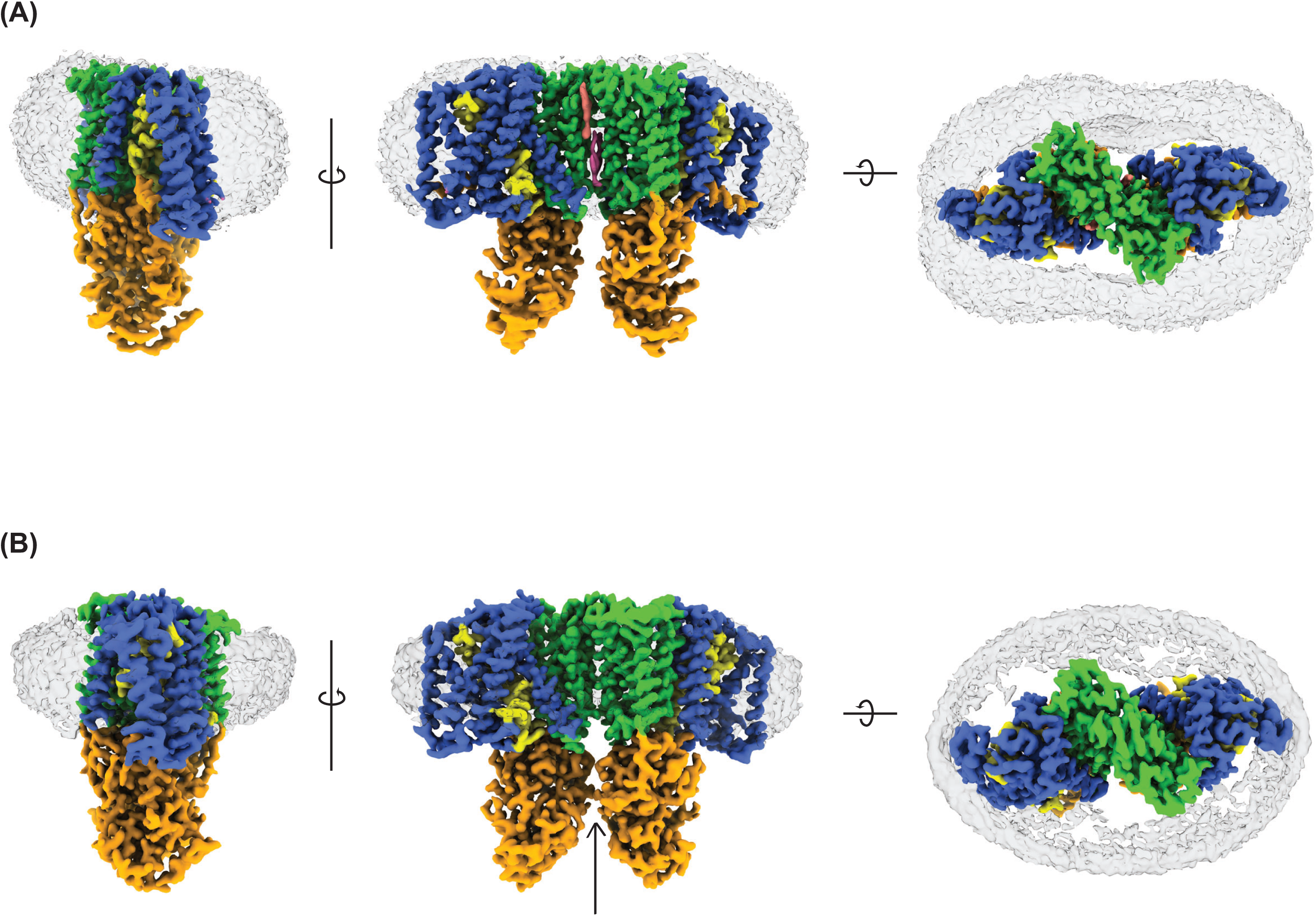
CryoEM maps of PaMprF. (A & B) Different views of the cryoEM maps of PaMprF in GDN micelle and nanodisc respectively. The central panel show the front view and the dimer interface. On the left, the side view of the enzyme and on the right, the view from the periplasm are shown. The arrow in panel B shows the proximity of the soluble domain and the compact nature of the enzyme in nanodisc. The cryoEM maps are coloured based on the functional regions of the enzyme and the model was used to define the boundary. The peripheral helices, dimer interface and re-entrant loops in the TMD are coloured in blue, green and yellow respectively. Soluble domain is coloured in orange. The detergent micelle and the nanodisc/lipid are shown as grey transparent surface around the TMD. The interface lipid and detergent at the dimer interface are shown in pink and salmon. In the periplasmic view, the continuous diffuse density in the GDN map and contrasting nanodisc belt can be clearly seen. The maps for the figures were generated using EMReady (He et al., 2023). To show the detergent/lipid belt, a second map with the macromolecule removed is overlaid.

**Table 1:**
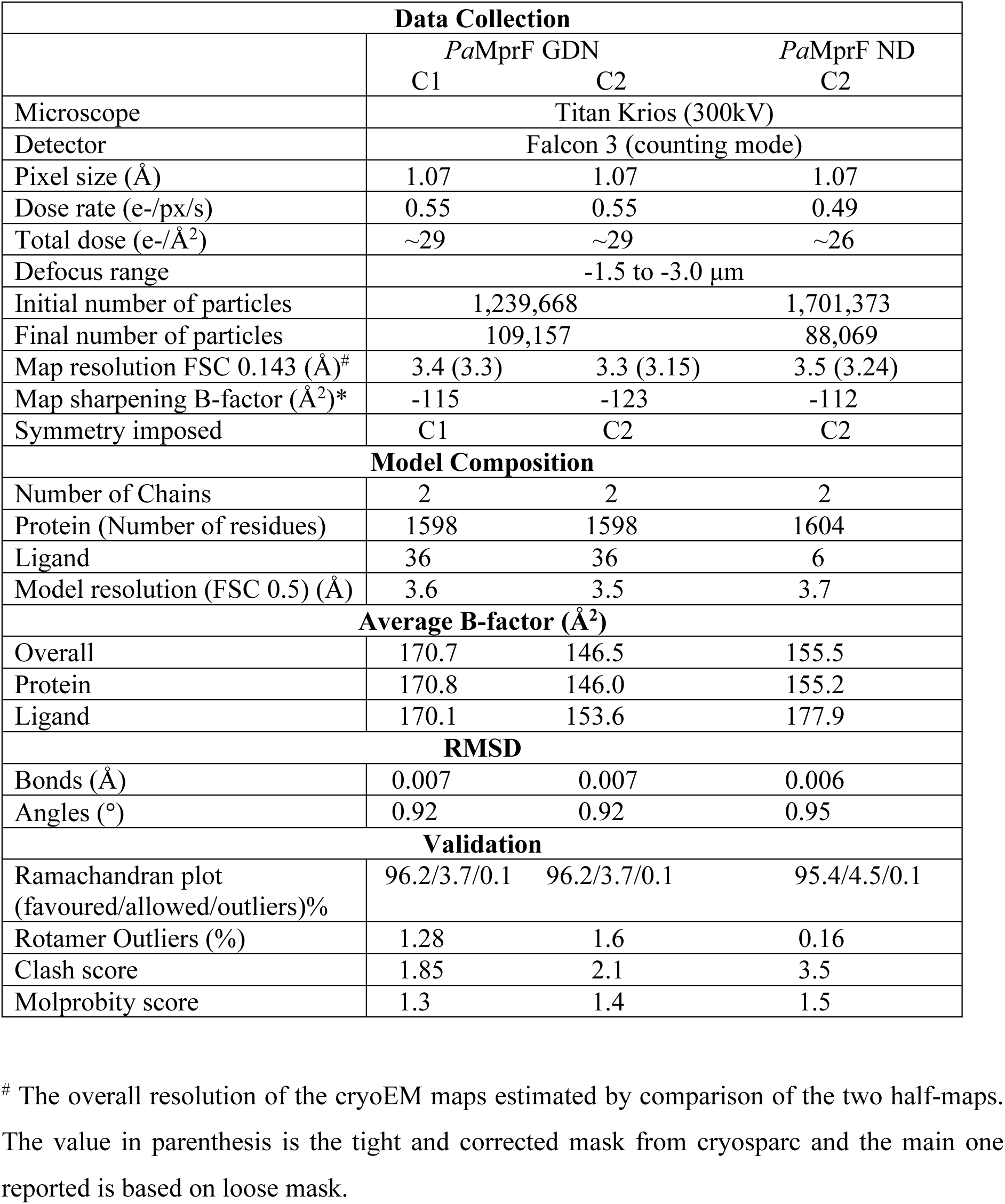
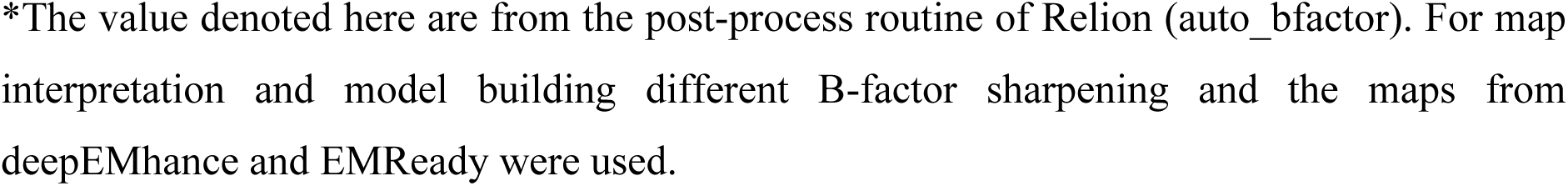
CryoEM Data collection and Model statistics.

| **Data Collection** |  |  |  |
| --- | --- | --- | --- |
|  | *PaMprF* GDN |  | *PaMprF* ND |
|  | C1 | C2 | C2 |
| Microscope | Titan Krios (300kV) |  |  |
| Detector | Falcon 3 (counting mode) |  |  |
| Pixel size (Å) | 1.07 | 1.07 | 1.07 |
| Dose rate (e-/px/s) | 0.55 | 0.55 | 0.49 |
| Total dose (e-/Å<sup>2</sup>) | ~29 | ~29 | ~26 |
| Defocus range | -1.5 to -3.0 μm |  |  |
| Initial number of particles | 1,239,668 |  | 1,701,373 |
| Final number of particles | 109,157 |  | 88,069 |
| Map resolution FSC 0.143 (Å)<sup>#</sup> | 3.4 (3.3) | 3.3 (3.15) | 3.5 (3.24) |
| Map sharpening B-factor (Å<sup>2</sup>)* | -115 | -123 | -112 |
| Symmetry imposed | C1 | C2 | C2 |
| **Model Composition** |  |  |  |
| Number of Chains | 2 | 2 | 2 |
| Protein (Number of residues) | 1598 | 1598 | 1604 |
| Ligand | 36 | 36 | 6 |
| Model resolution (FSC 0.5) (Å) | 3.6 | 3.5 | 3.7 |
| **Average B-factor (Å<sup>2</sup>)** |  |  |  |
| Overall | 170.7 | 146.5 | 155.5 |
| Protein | 170.8 | 146.0 | 155.2 |
| Ligand | 170.1 | 153.6 | 177.9 |
| **RMSD** |  |  |  |
| Bonds (Å) | 0.007 | 0.007 | 0.006 |
| Angles (°) | 0.92 | 0.92 | 0.95 |
| **Validation** |  |  |  |
| Ramachandran plot (favoured/allowed/outliers)% | 96.2/3.7/0.1 | 96.2/3.7/0.1 | 95.4/4.5/0.1 |
| Rotamer Outliers (%) | 1.28 | 1.6 | 0.16 |
| Clash score | 1.85 | 2.1 | 3.5 |
| Molprobity score | 1.3 | 1.4 | 1.5 |

<sup>#</sup> The overall resolution of the cryoEM maps estimated by comparison of the two half-maps. The value in parenthesis is the tight and corrected mask from cryosparc and the main one reported is based on loose mask.
\*The value denoted here are from the post-process routine of Relion (auto\_bfactor). For map interpretation and model building different B-factor sharpening and the maps from deepEMhance and EMReady were used.

To rule out the possibility of the CPD and its cleavage (by induction with IP_6_) causing any inadvertent effect, a 3D map of PaMprF was also obtained from a different construct where the cysteine protease domain was replaced by a precision protease site. This showed an identical architecture indicating that the flexibility of the synthase domain is an inherent nature of the enzyme in detergent micelle (Figure S2). This further prompted us to reconstitute the enzyme into MSP1E3D1 nanodisc with *E. coli* polar lipids and a cryoEM map with C2 symmetry at an overall resolution of 3.5 Å was obtained with slight anisotropy (Figure S3E and S4B, Table 1). The belt like feature of the membrane scaffolding protein (MSP) is clearly visible in the map (transparent grey surface in Figure 1B). The enzyme in nanodisc is more compact and the soluble domain is better resolved than the enzyme in GDN micelle (Figure 2A and B). This is also supported by asymmetric reconstruction, which shows no large difference (Figure S3 D). When the maps of the dimer from GDN and nanodisc are compared, it is apparent that the axis of one monomer appears to be tilted with respect to the other monomer (Figure 2C). Nanodiscs are primarily assumed to mimic the native lipid environment, but they are also known to constrict the transmembrane (TM) helices depending on the size of the disc formed (Dalal et al, 2024). We observe a similar compaction with PaMprF in MSP1E3D1 nanodisc and these changes in the TM regions can also be translated to the soluble domains.

**Figure 2:**
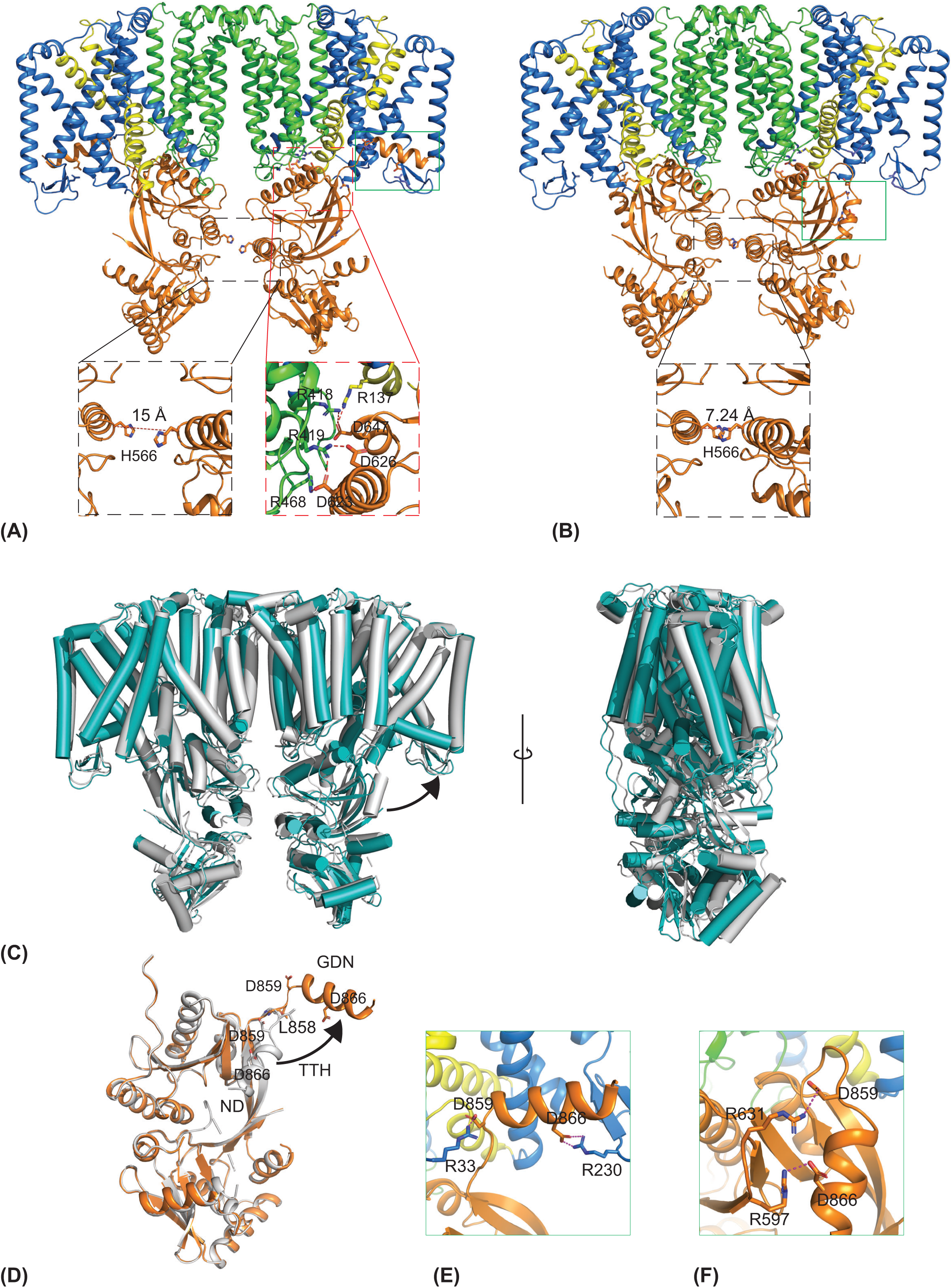
Models of PaMprF. (A and B) – models of PaMprF in GDN micelle and nanodisc respectively. The different regions of the molecule are coloured as in figure 1. The inset (black dashed line) shows the distance between the soluble domain in GDN and nanodisc as measured between the Cα of H566 residue. In addition, the electrostatic interaction at the interface of soluble and TM domain (marked in red dashed line) between D623 and R468/R418, D647 and R137/R418 and D626 and R419 are highlighted. (C) Comparison of model of PaMprF purified in detergent (teal) and nanodisc (gray). The tilt of the one of the monomers in nanodisc model is apparent. (D) Overlay of the soluble domains of PaMprF in GDN (orange) and nanodisc (gray) respectively. The position of the terminal helix towards the membrane is indicated by the black arrow. (E and F) The interaction of TTH in GDN and nanodisc. The residues D859 and D866 from TTH interact with the R33 and R230 respectively in the model derived from GDN EM maps. The same residues in the TTH of the nanodisc model interact with R597 and R631 in the soluble domain. The position of the TTH is marked with a green rectangle in the models shown in panel A.

### Structure of PaMprF

The model of the soluble synthase domain (PDB – 4v35) of *P. aeruginosa* was docked into the cryoEM maps and the TM helices initially built as poly-alanine model followed by assignment of the sequence. The fit of the models with density of both the GDN and nanodisc is shown in figure S4. The model was built prior to AlphaFold publication (Jumper *et al*, 2021) and the availability of the predicted structure. A comparison of the experimentally determined and the AlphaFold predicted structure reveals major deviation in the TM helices but less so in the soluble domain (Figure S5 A). There are a total of 14 transmembrane helices (TM) with two – TM 3 and 7 forming a pair of re - entrant helices facing each other at the centre of the membrane (Figure 2A and Figure S5 B). The monomers from both GDN and nanodisc align well with a rmsd of ∼0.3 Å for the TMD and some major differences in the soluble domain (discussed below). Comparison of the complete dimer model GDN micelle and nanodisc reveals an overall rmsd of ∼3.3 Å (Cα atoms) and a shift of 6.7 Å between the centroids of the reference monomer (Figure 2B). This can be appreciated with the proximity of the soluble domain in both the structures as measured between the residue H566 in the monomers (Figure 2A). As expected, the volume of the cavity comprising the dimer interface is larger 13089.1 Å^3^ in GDN micelle as opposed to 8434 Å^3^ in nanodisc.

The dimeric interface is formed by TM 11 and 12, ECH4 and lipid molecules [modelled as phosphatidylglycerol (PG)] and a detergent molecule fills the interface (discussed below). The synthase domain is connected to the TM domain through a loop between the last TM 14 and the first helix of the first GNAT1 fold (Figure S5 B). Of the two GNAT folds in the synthase domain, one closer to the membrane is better ordered. There are potential electrostatic interactions between the synthase and the TM domain (Figure 2A), two of these are formed at the helix 3 of the GNAT1 fold by D623 and D626 with R419 in TM10 and D623 with R468 in the loop between TM 12/13. The third one is formed by the re-entrant helix 3b (R137) and R418 with D647 on the fourth helix of GNAT1 fold (Figure 2A). Several regions of the GNAT2 fold are poorly resolved or disordered in both maps (GDN and nanodisc) and these include the residues between G806 and N832 comprising the 5^th^ α-helix, and the loop between the 2^nd^ and 3^rd^ α-helix. In the cryoEM maps, there are unmodelled densities in the soluble domain as well as at the interface between the membrane and soluble domain. These include an additional tubular density in PaMprF in GDN, which could be the 5^th^ α-helix of the second GNAT2 fold and a density in PaMprF in nanodisc at a position similar to the TTH in GDN (Figure S5 C, D and Figure 3A).

**Figure 3.**
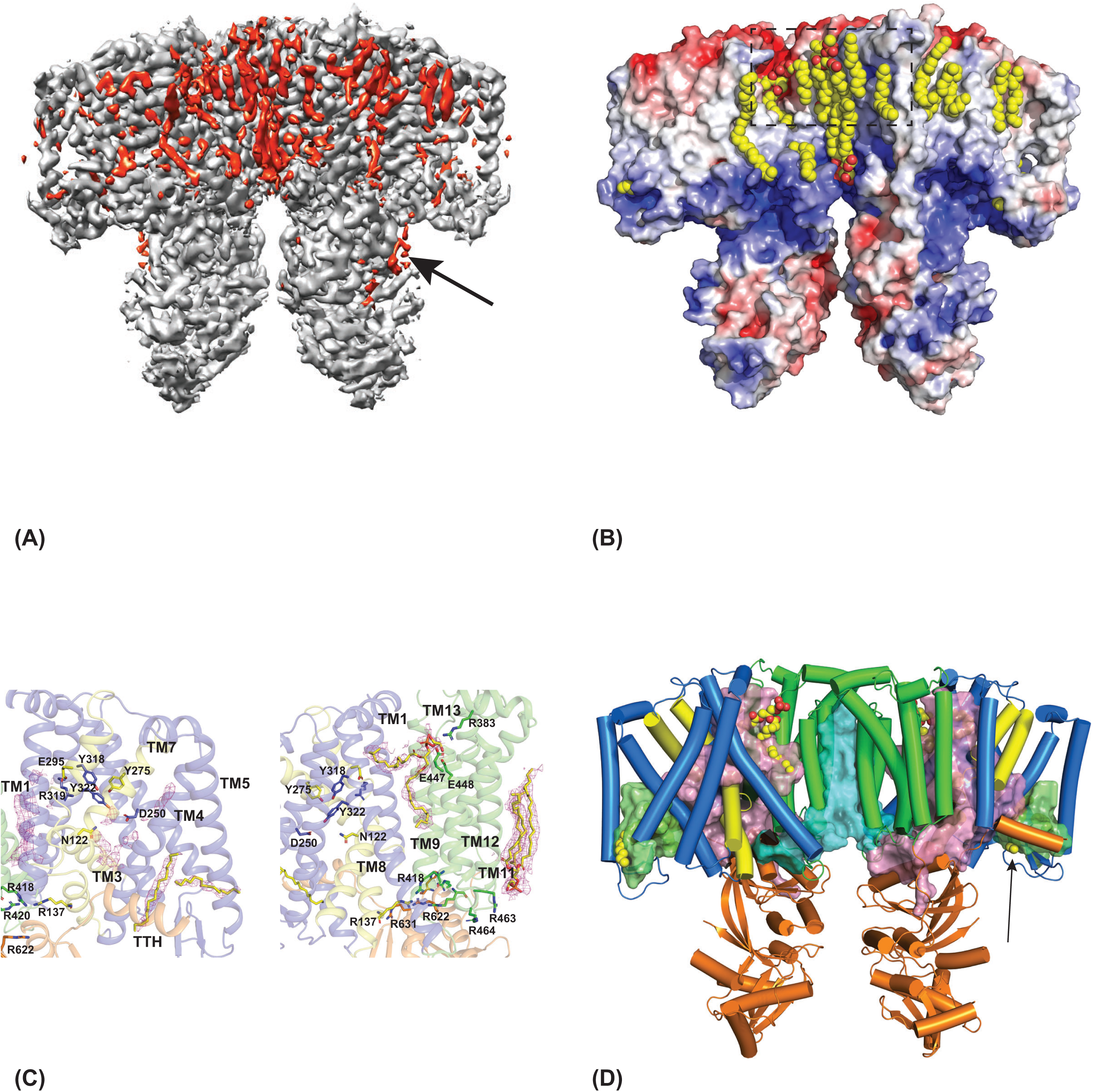
Lipids molecules in PaMprF in GDN micelle. (A) CryoEM map of PaMprF, showing unmodelled densities in red and the modelled protein density in grey. The lipid-like densities are clearly seen in the membrane domain and the fragmented density marked with an arrow is the unmodelled density in the soluble domain. The figure was made in Chimera with colour zone and split map option using the PaMprF model without any lipids included. (B) The electrostatic potential map of PaMprF is shown with lipids in sphere representation and coloured in yellow. The blue colour in surface map represents positive charge and red represents negative. The lipid molecules bind to the grooves and crevices of the enzyme. The lipid molecules that are lined up one after the other at the periplasmic side is marked with dashed black rectangle. (C) Some salient lipid molecules in stick representation (carbon atom coloured in yellow) are shown in the membrane domain (two views). The sharpened cryoEM map (in magenta mesh) at 5 α and carved 3 Å around the atoms is shown around the lipid molecules. Key residues that might play a role in lipid translocation are shown in stick representation. There are two partially ordered lipids close to TM4 and TM5 and one of them close to TTH, whose head group is likely to be close to the soluble domain. There are fragmented densities on either side of the re-entrant helices and are currently unmodelled. One PG molecule is modelled close to the periplasmic side and residues E447, E448 and R383, might mark the path for entry and exit of the lipids. Cluster of positively charged residues (R137, R418, R420, R463, R464, R622, and R631) line the interface between the membrane and soluble domain and are likely to play in interacting with lipid and or tRNA. (D The two cavities coloured pink and green that form the passage for the possible transport of lipids in the transmembrane domain. The smaller cavity coloured green is formed by the peripheral helices and the TTH, while the larger one coloured pink is present in the centre of the transmembrane domain, bifurcated by the re-entrant helices (coloured yellow) into two sub cavities. The cavity formed by the dimer interface is coloured cyan. Lipid molecules (sphere representation) are found in the cavity and the one close to TTH is marked with an arrow.

The last helix of the GNAT2 fold of the soluble domain, which we call as terminal transferase helix-TTH (the soluble domain is called synthase but its major role is in the transfer of aminoacid from tRNA to a lipid head group, so can also be termed transferase) occupies different positions in the maps obtained with GDN and nanodisc. In the model derived from the GDN map, the TTH is held in a parallel orientation to the membrane via the hydrophobic interactions of the non-polar residues, as well as electrostatic interaction (Figure 2C and 2D). Residues D859 and D866 in the TTH interact with R33 in TM1 and R230 in the β-sheet connecting peripheral helices TM 5 and 6 respectively (Figure 2D). In contrast, in the model derived from the nanodisc map, the TTH adopts a similar position to that observed in the crystal structure of PaMprF soluble domain alone, facing towards the solvent and interacting with the first β-sheet of the second GNAT fold (Figure S5D). In this conformation also, there are interactions of the acidic residues (D859 and D866) with the R597 and R631 (Figure 2E). An overlay of these models shows that the TTH moves as a rigid body with G857 acting as a pivot. The distance measured from the Cα distance of L858 between the two structures is ∼5 Å (Figure 2C). The different position of the TTH in the two maps is a salient feature and might play a role in the mechanism.

### Lipid molecules in PaMprF

A number of non-protein densities are observed in the cryoEM maps derived from the GDN micelle, some of them with classical pose of lipids with two acyl lengths and others that reveal short tube-like density, have been modelled as single acyl chains of either PG or phosphatidyl ethanolamine (PE) (Figure 3 A and B). These densities attributed as lipid molecules were also verified using a fofc omit map with servalcat (Yamashita *et al*, 2021). The number of lipid-like densities in the outer leaflet are greater in number perhaps indicating dynamic nature at the cytoplasmic leaflet (Figure 3B). Series of single acyl chain density is evident from one of the openings on the periplasmic side (marked in rectangle in figure 3B) and in the surface representation, it is evident that many of these lipid molecules fit into the grooves of the enzyme. Among the modelled lipid molecules, we would like to describe a few that might be of functional relevance (Figure 3C). Two partially ordered lipids molecules are found close to TM4/TM5 and there are fragmented densities at either side of the re-entrant helices TM3 and TM7 (Figure 3C), which have not been modelled currently. But these unmodelled densities are close to key residues that might play a role in lipid translocation (Song et al 2021). Of interest in the PaMprF structures are the presence of cluster of charged residues including R137, R418, R419, R420, R463, R464, R468, R622, R631, and H629. These residues are positioned at the interface of the TMD and the synthase domain and could play roles in interacting with acidic residues, (Figure 2A) to lipid head groups and importantly in their interaction with tRNA at the membrane interface. At the dimer interface, there is density for two lipid molecules at the inner leaflet, one completely ordered and the other partially ordered and are currently modelled as PG molecules but could also be cardiolipin (Figure 3B and 4G). R463 and R464 can form potential hydrogen bond interactions with lipids from the same as well as the opposite monomer. In addition, a lipid is modelled close to the periplasmic side, which is surrounded by TM1, TM8, TM9 and TM13 (Figure 3C) and opens towards the periplasmic space. An equivalent density at the same position is also observed in the nanodisc map and it could be Ala-PG but we do not have structural or biochemical evidence at the moment to support this and hence modelled as PG. This lipid is in close proximity to residues R383, E447 and E448 (Figure 3C).

**Figure 4.**
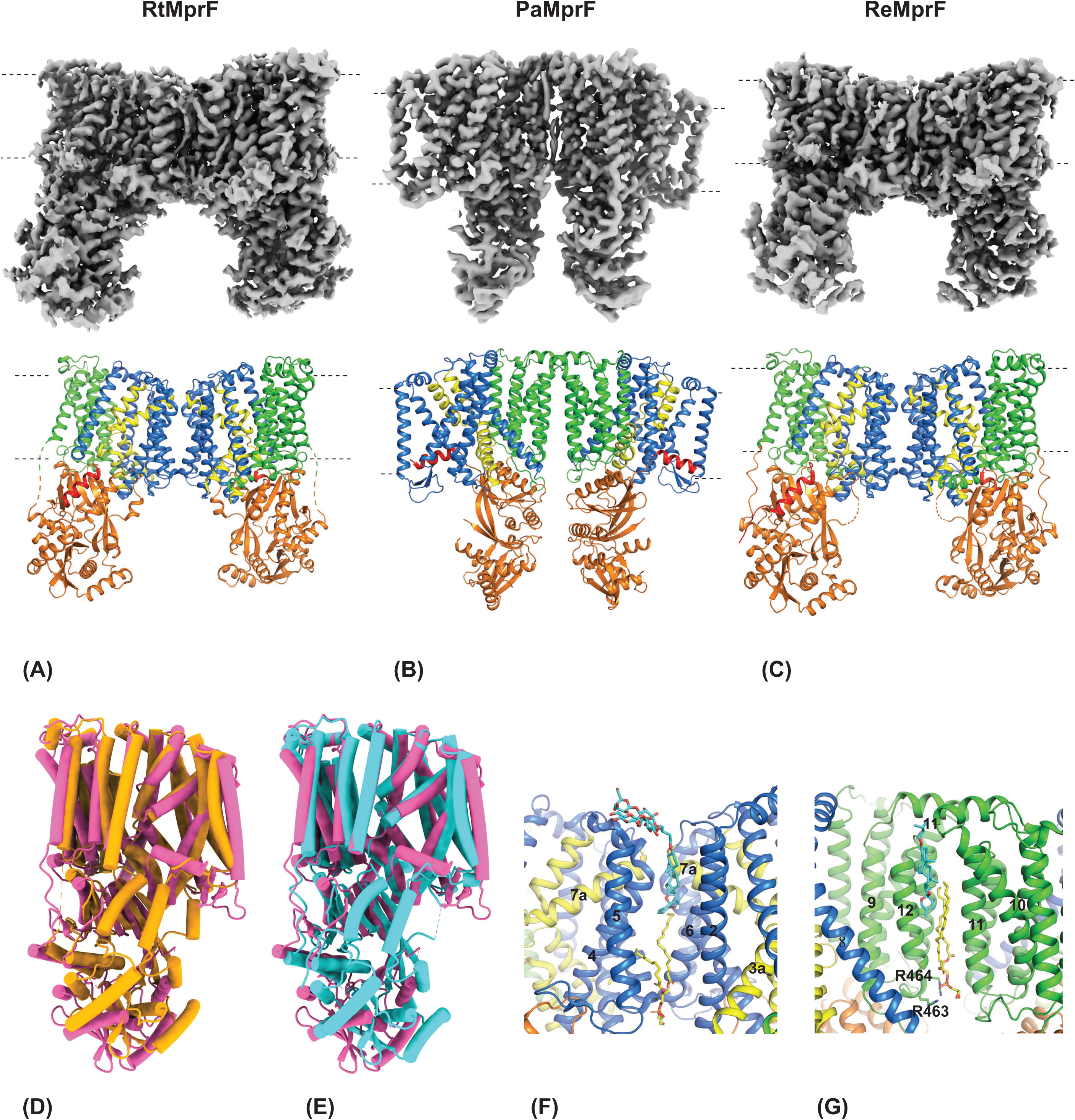
Comparison of MprF structures. (A-C) Comparison of dimeric interface from different MprF enzymes. The maps (top) and the models (bottom) of MprF from *Rhizobium tropici*, EMD-30869 and PDB–7DUW (left), *Pseudomonas aeruginosa* (center) and *Rhizobium etli*, EMD-31445 and PDB–7F47 (right) are shown. The membrane boundary is marked by dashed line. The models are coloured as in figure 2 with the N-terminus of the TM region in blue, re-entrant helices in yellow and C terminal TM helices in green. The soluble domain is coloured in orange. The difference in the dimer interface is clearly visualised. (D) and (E) Overlay of PaMprF monomer (magenta) with monomers of RtMprF (orange), and ReMprF (cyan) showing larger deviation in the membrane domain. (F) and (G) show the dimer interface of RtMprF and PaMprF respectively. The detergent GDN is found in the same position in both RtMprF and PaMprF despite the dimer interface formed by different TM helices. In RtMprF, TMH 4, 5 and 6 form the dimer interface but TMH 10 and 11 in PaMprF.

Analysis of the PaMprF enzyme for cavities reveals interesting features beyond the one observed at the dimer interface in both GDN and the nanodisc maps. The arrangement of the TM helices in PaMprF gives rise to a larger cavity encompassing the TM domains centred around the re-entrant helix (coloured pink in Figure 3D). Due to the movement of TTH in the GDN maps, this cavity has slightly smaller volume (5365.1Å^3^) than in nanodisc (6017.7 Å^3^). This larger cavity can be divided into a sub-cavity formed by TM helices 4, 5, 6 and the β-strands connecting TM 5 and 6 and the TTH (coloured green in Figure 3D), which is about 974 Å^3^ in volume and harbours a partially ordered lipid molecule perpendicular to the TTH, and co-planer to the membrane surface (Figure 3C). This cavity perhaps provides a path for holding lipids before or after the transfer of amino acid from tRNA. Taken together with partially or fully ordered lipids in the PaMprF in particular those lined up towards the periplasmic side or the outer leaflet (Figure 3B and 3D), and the vicinity to the cavities described above indicates the possible path taken by the lipids to enter and exit the enzyme.

### Comparison of MprF structures

Two structures of MprF from *Rhizobium tropici* and the map as well as the model for *R. etli* are now available (PDB-7F47 and EMD-31445) (Song *et al*, 2021). The EM maps from *R. tropici* were obtained in nanodisc with the enzyme purified either in DDM or GDN (but initial solubilisation in each preparation was performed with DDM). From the first glance, it was evident that the maps of *P. aeruginosa* and *Rhizobium sp.* are different (Figure 4 A), obvious from the position of the soluble domain with respect to the TM domain as a result of different dimer interfaces. The monomeric models of these enzymes align with an overall RMSD for Cα atoms of 3.3 Å and 3.1 Å for RtMprF and ReMprF respectively (Figure 4D and E). Not surprisingly, the interface and the salt bridges between the soluble and TM domains are also conserved in RtMprF. Greater deviation is observed in the TM domain of these structures (2.1 Å and 1.7 Å for RtMprF and ReMprF respectively) when compared to PaMprF in GDN micelle, while the rmsd for the soluble domain alone is ∼1 Å. The location of the helix corresponding to TTH in the structures of Rhizobium species are similar to PaMprF (nanodisc) model, i.e., away from the membrane interacting with the soluble domain. In PaMprF, TM helices 11 and 12 and external helix (ECH4) (coloured green in Figure 4A) form the dimer interface, while in both the Rhizobium MprFs, this is made of TM helices 4, 5 & 6 (coloured blue in Figure 4B). Perhaps, an interesting feature is the position of the ordered lipid and detergent, which is conserved in both the PaMprF and RtMprF maps despite the difference in the dimer interface (Fig 4G and F). Thus, the difference in the oligomeric arrangement of MprF from different homologs is intriguing and raises the question which of these arrangements is biologically relevant or this is perhaps an effect arising from the choice of the detergent.

### Effect of Detergent on PaMprF

To address if PaMprF dimerization interface is real or an effect of detergent, we employed two different approaches – screening and analysing the effect of detergents in combination with Blue Native (BN)-PAGE and chemical cross-linking in the membrane. In the initial stages of the PaMprF project, we had performed a detergent screen with various detergents and lipids both during solubilization and purification. Solubilisation of the PaMprF from the membranes with N-dodecyl β-D-maltoside (DDM) and exchange to other detergents during purification rendered the enzyme unstable (Figure 5A). However, solubilisation with LMNG or digitonin and subsequent exchange to GDN provided the best profile in the BN-PAGE gel (as well in the size-exclusion chromatography). On hindsight, the importance of GDN can be appreciated by its localisation at the dimer interface (Figure 4G). It was evident from BN-PAGE gels that predominantly monomeric bands were observed when solubilization was performed with DDM and subsequently exchanged to GDN (Figure 5B). In contrast, when LMNG is used for solubilisation and exchanged to GDN, a strong dimeric and some higher molecular weight bands were also observed (Figure 5B). It was thus evident that the first step of extraction from the membrane with the milder detergents such as LMNG or digitonin is essential for the stability of PaMprF.

**Figure 5.**
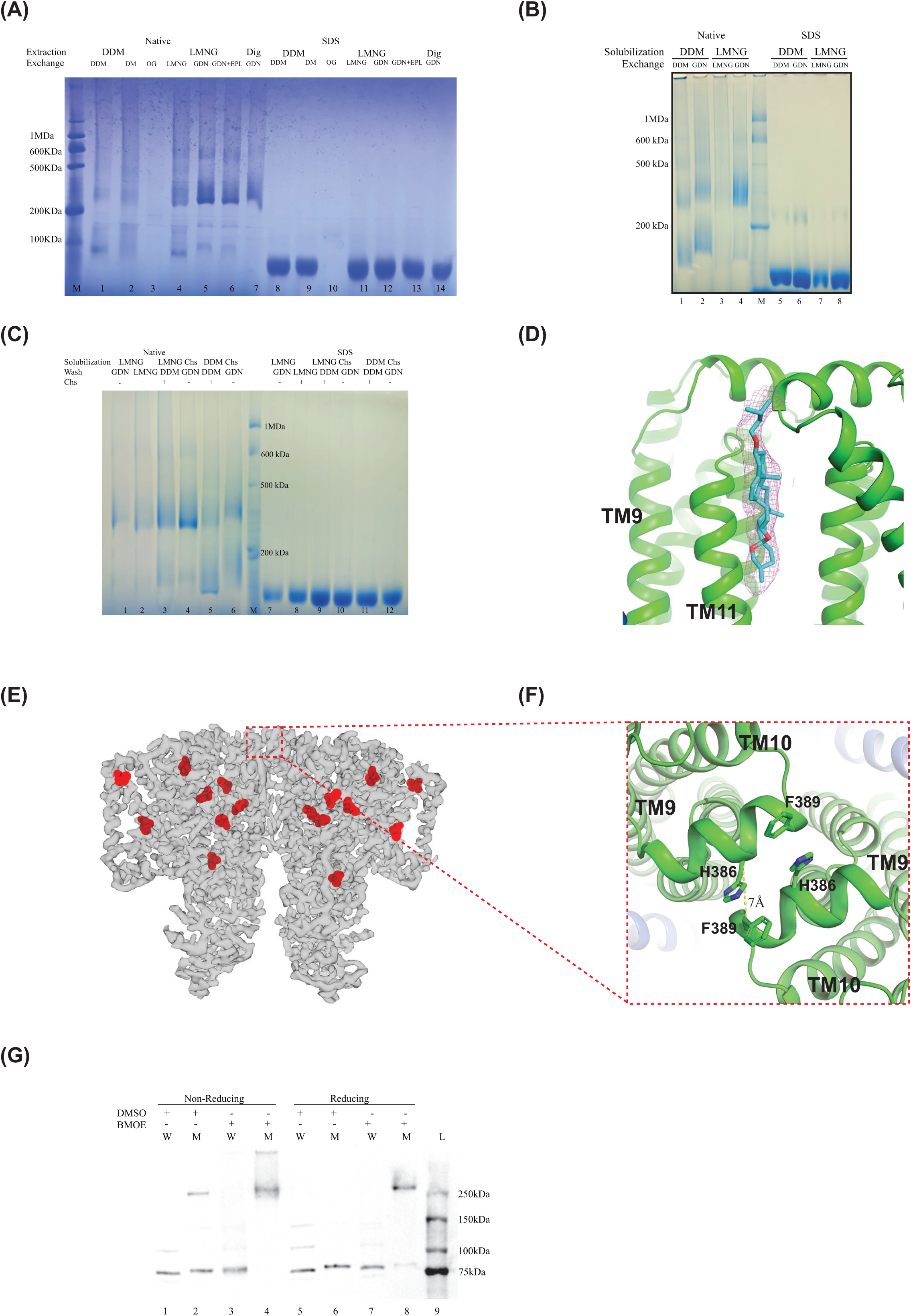
Effect of detergents on PaMprF and oligomeric state in the membrane. (A) Detergent screen of PaMprF assessed by BN PAGE showing native samples immediately after the marker lane (M) followed by SDS treated samples as control, where the oligomers dissociate into monomers. In this experiment, solubilisation was performed either with DDM (lanes 1-3), LMNG (lanes 4-6) or digitonin (lane 7) and the detergent used for purification was exchanged on the column. Detergent used for purification (from lane 1-7) are DDM, DM, OG, LMNG, GDN, GDN with *E. coli* polar lipids (EPL), and GDN, respectively. Lanes 8-14 are equivalent to lanes 1-7 but treated with 1% SDS showing monomeric band except for OG, where no clear band was observed. 10 μg of purified protein was loaded in each lane. (B) BN PAGE showing the comparison of DDM or LMNG extraction on PaMprF. Lanes 1-4 and 5-8 are native and SDS denatured samples respectively. Note the difference in the profile of lane 2 and 4 indicating the dissociation of DDM solubilised PaMprF, even when exchanged to GDN. Approximately 15 μg PaMprF has been loaded in each lane. (C) BN PAGE showing the effect of cholesterol hemi succinate (CHS) during extraction and purification on the stability of PaMprF. On-bead cleavage was performed using equal volumes of IP_6_ buffer and 20 µl was loaded in each lane. The native samples in lanes 1-6 and their corresponding SDS denatured samples (lanes 7-12) indicate the varying stability and yield under different conditions respectively. Note the band intensity of LMNG CHS solubilised PaMprF in lane 10, compared to the one without CHS in lane 7 (CHS – cholesterol hemi succinate). The marker in all the 3 BN PAGE gels shown here is solubilised mitochondrial fraction in DDM. (D) The cholesterol skeleton of GDN detergent modelled in the density present between the two monomers towards the upper leaflet. The density of the GDN (carbon atoms in cyan) is shown in a magenta mesh using the sharpened map of PaMprF (GDN) with map double option in pymol and contoured at 4 σ. (E) cryo-EM map (light gray) of PaMprF with the position of the cysteine residues shown in red. (F) Zoomed in view of the dimeric interface as seen from the periplasmic side showing the loop between TM helices 9 and 10. Residues H386 and F389 are shown in stick representation. C*α*-C*α* distance is ∼7Å. (G) Western blot showing crosslinking of PaMprF with bismaleimidoethane (BMOE). Lanes marked W and M are wild type and mutant respectively. Lanes 5-8 are samples with 20 mM DTT added. Lane 2 shows that the H386/ F389 mutants has the propensity to form dimer even without the addition of a crosslinker and when BMOE is added, no monomers are observed (lane 4). This indicates the proximity of the residues in the membrane. Protein ladder (Biorad Cat No. 1610375) is loaded in lane 9. Introduction of these additional cysteines results in slower migration of the monomer band of the mutant (lanes 2, 6) and aberrant migration of the dimer at >250 kDa (expected size of the cross-linked dimer is ∼150 kDa).

As a consistent density of cholesterol skeleton from GDN was observed at the dimer interface in the EM maps of PaMprF from different biochemical preparations and also observed in the cryoEM map of RtMprF (Figure 5 D), we asked if cholesterol hemi succinate (CHS) (commonly used in eukaryotic membrane protein purification) can be added during solubilisation with LMNG or DDM to stabilise PaMprF. It was observed that the presence of CHS during the solubilization step improves the yield of PaMprF, but does not remove the necessity of GDN during purification (Figure 5C).

Detergents are known to induce artefacts in the structure of membrane proteins (Ravera *et al*, 2022; Wu *et al*, 2020; Tate, 2006). Although, the nanodisc structure of PaMprF with lipids indicate the same dimer interface, the initial steps in purification still involved the addition of detergent and it can thus be argued that the observed organisation could be a result of detergent induced artefact. To probe the organisation of the PaMprF monomers in the membrane, cysteine crosslinking was performed. The enzyme PaMprF has a total of 8 cysteine residues; seven of these are in the TM domain and one in the soluble domain (Figure 5E). The loop connecting the TM 9 and 10 on the periplasmic side forms a short helix, which is antiparallel to its counterpart on the other monomer at the dimer interface. H386 of one monomer is ∼7 Å away from F389 of the opposing monomer (the distance between the C*α* atoms and Figure 5F). Both these residues were mutated to cysteine (H386C/F389C) to determine the proximity of these helices in the membrane environment (note that the native cysteines have not been mutated). The membrane fractions of the WT and the H386C/F389C double mutant of PaMprF were analysed in non-reducing conditions with anti-strep antibody (Figure 5G). Even in the absence of cross-linker, a higher molecular weight band along with the monomer band of the mutant was observed. With addition of a cross-linker, only the higher molecular weight band was observed and this could be partly dissociated with the addition of DTT (Figure 5G). This higher molecular weight band is likely to be the dimeric form of PaMprF but shows aberrant migration in SDS-PAGE. Thus, the cross-linking experiment supports the dimer interface of PaMprF in the membranes as observed in the cryoEM maps obtained with the purified enzyme in GDN micelle and reconstituted in nanodisc.

## Discussion

Microbial resistance to small molecules such as antibiotics involved in translation inhibition can arise from small changes, for example methylation of rRNA bases (Osterman *et al*, 2020). Similarly, the resistance to CAMPs widely produced by many organisms is achieved by the transfer of an amino acid from tRNA to a lipid head group, a relatively simple solution using the abundant charged tRNAs in the cells. Mechanistically, this is also interesting as the protein that performs this function requires both a charged surface and a hydrophobic region for tRNA and a lipid molecule respectively to bind and bring the head group of lipids in close proximity to tRNA for the amino acid to be transferred. Additionally, the lipid with the modified head group needs to be translocated from one leaflet to another. Such dual functional enzymes are more commonly found in pathogenic bacteria conferring resistance to CAMPs and several enzymes from multiple species have been characterised (Fields & Roy, 2018; Slavetinsky *et al*, 2012).

In this study, we report the cryo-EM structures of the PaMprF both in GDN micelle and MSP nanodisc. The maps reveal a dimeric enzyme as observed recently in the homologous Rhizobium enzymes (Song *et al*, 2021). While they share the same monomeric fold but have different dimeric interface. In PaMprF, the dimeric interface is mediated by TM 11 and 12 and in Rhizobium enzymes, this interface is formed by TM 5 and TM 6 (Figure 4A). The origin of this difference is intriguing and might represent different states of the enzymes or the effect of detergents used in the process of extraction and purification. In the study of membrane proteins, detergents are indispensable, and occasionally, they are known to disrupt the native conformation of membrane proteins (Ravera *et al*, 2022; Wu *et al*, 2020; Tate, 2006; Warne *et al*, 2008; Chipot *et al*, 2018). DDM is one of the most widely used detergent in membrane protein biochemistry and structural biology and in recent times, milder detergents such as LMNG, digitonin and its synthetic analogue GDN with a cholesterol backbone have been used to study a number of labile membrane proteins (Ehsan *et al*, 2022; Lee *et al*, 2020; Chae *et al*, 2010). When PaMprF was screened with detergents in different combinations and analysed by BN-PAGE, it was evident that the use of DDM for initial solubilisation made the enzyme less stable and LMNG was required (Figure 5A). Subsequent exchange to GDN kept the enzyme stable only if extracted initially with LMNG or digitonin. While cholesterol or cholesterol-like molecules are rarely found in prokaryotes (Lee et al, 2023), still cholesterol hemi succinate has been used to purify membrane protein from prokaryotes (Shen et al, 2023). For PaMprF, the use of CHS with LMNG improved the yield but not the stability (Figure 5C). The importance of GDN can be appreciated by its presence at the dimer interface, where it is sandwiched between TM helices 11 and 12 in PaMprF. Surprisingly, GDN along with a PG molecule occupy similar positions in both PaMprF and RtMprF despite the difference in the dimer interface (Figure 4G).

When GDN purified PaMprF was reconstituted into nanodisc, the dimeric assembly similar to that in the micelle was observed (Figure 1 and 2). Although, the two structures of RtMprF were also determined in nanodisc, the initial solubilisation was performed with DDM, followed by either purification in DDM or GDN, both these data sets show fuzzy density for soluble domain in 2D classes indicating heterogeneity (Song et al, 2021). With PaMprF, when DDM was used for solubilisation low yield of enzyme was obtained and stability was compromised (Figure 5A, B and C). Further, we introduced two cysteine residues at the dimeric interface of PaMprF and a homo bi-functional cross-linker BMOE was used to cross-link the enzyme in the membrane. The cross-linking experiments clearly show that the dimeric architecture of PaMprF observed in micelle is also present in the membrane. Thus, we conclude that MprF architecture/oligomeric state is sensitive to the type of detergent used and the arrangement of monomers as observed in PaMprF is realistic.

The TM domain of MprF has been postulated to perform the flipping of lipid from the inner to the outer leaflet and they are characterised by 14 TM helices and re-entrant helices commonly found in transporters (Drew & Boudker, 2016). Two cavities are observed on either side of the re-entrant helices, and in both the PaMprF GDN and nanodisc maps, non-protein densities attributed to lipids are observed (Figure 3A and 3D). In the region close to the outer leaflet of the enzyme, series of lipid densities are found and they form a continuous array from the re-entrant helices indicating the path taken by the lipids (Figure 3B).

The synthase domain of MprF has two GNAT folds and is well studied structurally and mutagenesis of conserved residues has indicated that they play a role in specificity and interaction and the substrates themselves play a role in catalysis (Hebecker *et al*, 2015, 2011). In PaMprF GDN map, synthase domains reveal greater flexibility but are more compact in the nanodisc map (Figure 2A). The models of the synthase domain derived from EM maps are largely identical to the structure of the isolated soluble domain with few crucial differences. The TTH is buried in the membrane, perpendicular to the TM domain in the GDN map but exposed to the solvent in the nanodisc structure (Figure 2C). The TTH is an amphipathic helix and has been predicted to be a putative TM helix as well (Roy & Ibba, 2009). Deletion of this helix in PaMprF is known to abolish AlaPG synthesis (Hebecker et al, 2011). The two structures of PaMprF from detergent and nanodisc perhaps reveal two different conformations of TTH, which is at the interface of the membrane or placed away from the membrane perpendicular to the membrane surface. Two acidic residues D859 and D866 from TTH in both the structures form electrostatic interaction with arginine residues with the TM as well as the soluble domain (Figure 2 E and F). This switch of TTH between the two positions is a possible mechanism to allow tRNA molecule to bind and for the lipids to come closer to tRNA.

Previous structural study of the synthase domain (*Bacillus licheniformis*) with lysine amide that mimics the amino acid at the end of tRNA and comparison with tRNA analogue bound FemX was used to extrapolate the tRNA binding site and conserved residues in the region, whose mutation resulted in reduction or loss of activity (Hebecker et al, 2015). In that report, a lipid binding site opposite to the tRNA binding site was postulated as a tunnel that could accommodate a PG molecule and place it closer to the lysine amide (Figure 6A and 6B). In the context of the full-length PaMprF, the lipid binding site is exposed to solvent and as previously observed in the RtMprF, there is no continuous tunnel or path from the membrane domain to the soluble domain that might protect the hydrophobic part of the lipid molecule. In addition, the compaction observed in the nanodisc structure will make the previously predicted PG binding site not easily accessible (Figure 2B).

**Figure 6:**
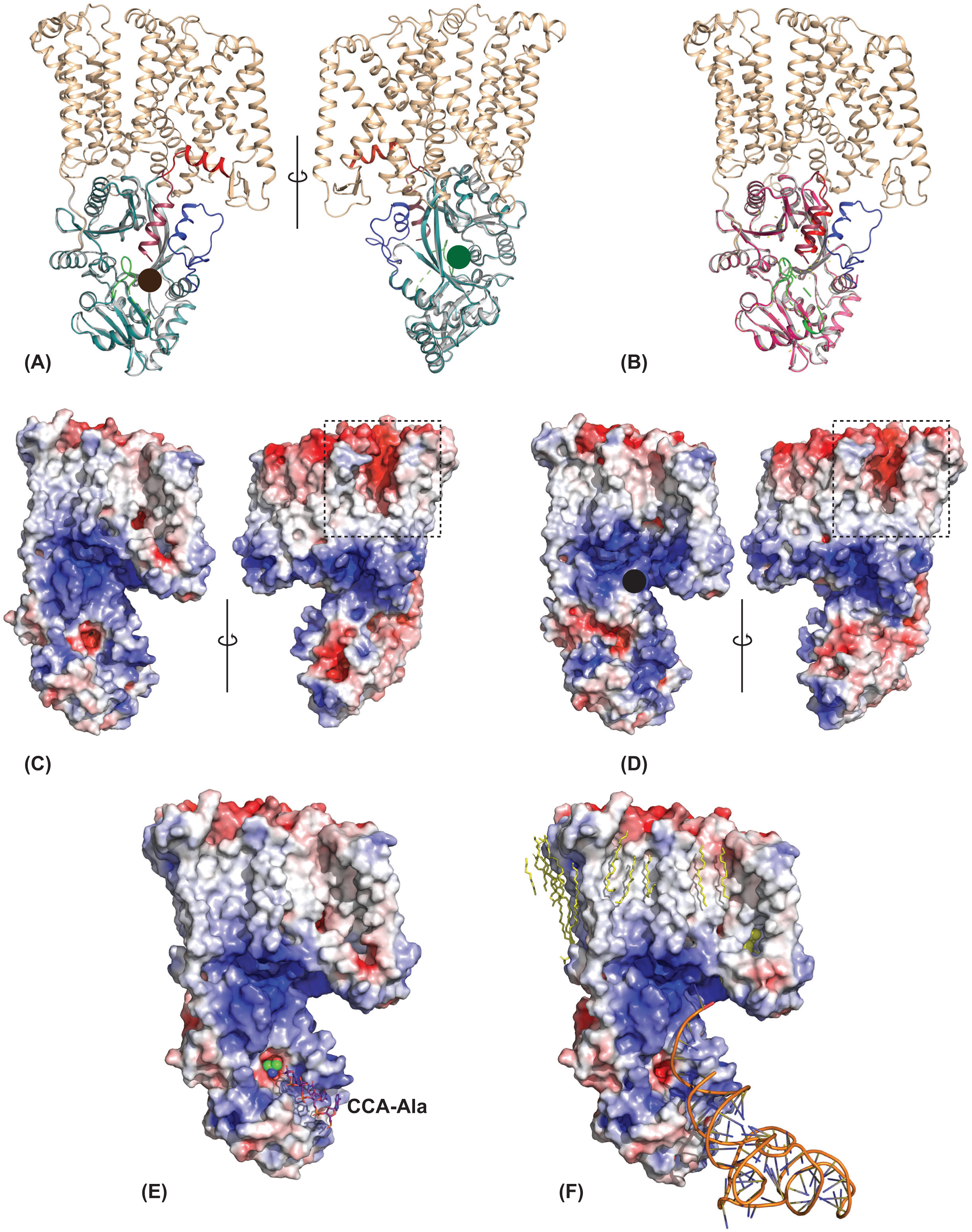
A model for tRNA binding and aminoacylation. (A) An overlay of the monomer model from the GDN cryoEM map and the soluble domain (4v35). In the cryoEM GDN derived model, the TM domain is coloured in wheat, TTH in red and the synthase (soluble) domain in teal. In the crystal structure of the soluble domain, the TTH is coloured in brown, the helix 6 of GNAT2 fold in blue and disordered loop in green, while the rest of the molecule is in gray. Two views of the molecule are shown. The cavities marked in brown and green circle are the postulated tRNA binding and the lipid binding sites, respectively (Hebecker *et al*, 2015). (B) An overlay of the models of the nanodisc structure and the crystal structure of the isolated soluble domain (4v35). The colouring of the crystal structure is the same as in panel A and the synthase domain of the nanodisc is in magenta and the TTH occupies a similar position. (C and D) Electrostatic surface representation of GDN and nanodisc monomer models. Two views are shown and coloured with −5 to +5 kT. The black dotted box shows the possible entry/exit site for the lipid at the periplasmic side and the outer leaflet (see also figure 3B). Note the general and continuous positive surface in the synthase domain and at the interface. In panel D, the black circle marks the position of TTH, which might shield the surface and prevent the binding of tRNA. (E) A simple docking exercise using CCA-ala in the predicted tRNA binding site (brown circle in panel A). The alanine residue fits snugly in the cavity within the synthase domain (also compare with panel C). However, there is no clear path for lipids to reach this site without exposing itself to the solvent. (F) A tRNA molecule positioned such that the CCA end with the amino acid is close to the positive surface at the cytoplasmic leaflet. The switch of TTH closer to the membrane exposes the charged residues and also forms a wall of the cavity at the far side of the enzyme. In this model, the lipid will still be buried with the TM domain (as shown here with partially ordered lipid in yellow sphere) and only the head group will be exposed to the solvent and come closer to the tRNA. For the figures in panels E and F, the model derived with GDN was used.

The electrostatic potential calculation of PaMprF show a positive surface that extends from the cytosolic side of the membrane domain and the soluble domain (Figure 6C and D). Extrapolating from the FemX bound partial tRNA analogue and UDP (Fonvielle et al, 2013), a groove in the synthase domain is observed that can hold the tRNA. However, the membrane domain is further away and there is no easy path for lipid to reach the tRNA binding site. We also realise that even if such path were formed by local rearrangement, there is no easy mechanism for the lipid to go back to the membrane domain after the amino acid transfer. An alternative and elegant solution that can be derived from the structures presented here is if the CCA end of the tRNA molecule is closer to the inner leaflet of the membrane. Supporting this possibility is the presence of positive patches that can hold the tRNA contributed by charged residues from both the synthase and TM domains (Figure 6C and D). Comparison of the synthase domain from different structures of PaMprF shows that for tRNA to bind close to the membrane domain, structural change is likely required. In the cryoEM maps, the flexible or disordered regions of the synthase domain include the loop connecting the 2^nd^ and 3^rd^ α-helix, helix 5 of the GNAT2 fold and the TTH. Of these, the movement of TTH as observed in the GDN cryoEM structure towards the membrane domain might have two roles. First, it allows for tRNA binding as the two arginine that interacted with aspartate in TTH are now free to interact with incoming tRNA and secondly, by exposing hydrophobic residues it forms part of the cavity, where lipid molecules can be housed before and after amino acid transfer from tRNA (Figure 3D and Figure 6E and F). Among the missing links in the current cryoEM maps is the lack of well-ordered density for lipid molecules on the inner leaflet closer to the re-entrant helices but it is reasonable to assume from the cluster of positive charge that there will be lipid molecules and are dynamic. The synthase domain with its positive patches can attract tRNA towards it and one possibility is that it can push the TTH and further interact with residues close to the cytoplasmic leaflet. The lipid molecule could be housed in the cavity (formed by TTH and TM 4, 5 and 6, coloured green in figure 3D) and the amino acid is transferred to the headgroup. Upon alanylation, the AlaPG translocates through the cavity formed by TMH 1, 7, 8, and 11, with the re – entrant helices facilitating the transport of charged head group of the AlaPG using ionic interactions formed by its polar residues present at the loop region in these helices. Towards the end of this process, the AlaPG aligns parallel to the lipids in the membrane surface and its head group is now facing diagonally opposite to the putative alanylation site.

The cleft acts as the final exit site for the lipids that appear out of the sub cavities on the other side of a monomer and are then laterally moved towards the other monomer into these clefts arranged in a parallel array (Figure 3B).

While this looks like an attractive prospect, residues involved in the mechanism cannot be deduced although several charged residues are in the vicinity. Two caveats in the present study include the lack of a structure with tRNA or its analogue and an *in-vitro* functional assay that can be used to measure both synthase and flippase activity simultaneously. Nevertheless, the distinct arrangement found between MprF from two homologues, structural changes in the GDN and nanodisc structures of PaMprF and the presence of lipids within the TM domain and on the periphery provide an avenue to explore the mechanism of these bi-functional enzymes.

## Methods

### Protein expression and preparation of membranes

MprF gene was amplified from the genomic DNA of *Pseudomonas aeruginosa* and cloned into pET22b vector carrying a StrepTagII at the N terminus and a Cysteine protease domain followed by hexahistidine tag at the C terminus. H386C/F389C double mutant was generated by a single pair of oligonucleotides using site directed mutagenesis (Costa et al, 1996). All the constructs were verified by sequencing. For expression, the plasmids were transformed into *E. coli* BL21 (DE3) C41 cells (Miroux & Walker, 1996). A single transformant colony was inoculated in LB media containing 100 μg/ml ampicillin and grown overnight at 37°C. This was used to inoculate larger cultures with 100 μg/ml ampicillin and allowed to grow at 37°C till the OD_600_ reached 0.5. The temperature was reduced to 18°C and induction was carried out by adding 1 mM IPTG for 16 hours. All the following procedures were performed at 4°C unless specified otherwise. The cells were harvested by centrifugation at 4000 g, resuspended in 100 mM Tris pH 8, 1 mM PMSF, 4 mM MgCl_2_, 0.5 mM CaCl_2_, 4 μg/ml DNase and 5mM ß-mercaptoethanol. Cells were lysed by passing the suspension thrice through Emulsiflex-c3 (Avestin) at 15,000 psi. The unbroken cells were removed by centrifugation at 15000 g for 30 minutes and supernatant was subjected to ultracentrifugation at 200,000 g in a Ti45 rotor (Beckmann Coulter) for 2 hours to obtain the membrane fraction. The membranes were homogenised and resuspended with 50 % wt/vol in a buffer A containing 50 mM Tris pH 8, 200 mM NaCl, 5 mM ß-mercaptoethanol and stored at −20°C. The total protein concentration in the membrane fraction was estimated using Bradford reagent (Bradford, 1976).

### MprF Purification in GDN

Membranes were thawed and diluted to a final protein concentration of 5 mg/ml using Buffer A containing 20 mM imidazole and solubilized using 1 % LMNG at 4°C for 1 hour and centrifuged at 150,000 g for 30 minutes to remove the insoluble fraction. The supernatant was then applied to a 5 ml HisTrap (GE Healthcare) column pre-equilibrated with buffer A containing 0.006% GDN (buffer B) and 20 mM imidazole. The column was washed with 30 CV of buffer B containing 40 mM imidazole. Full-length MprF was then eluted with buffer B supplemented with 200 mM imidazole. 200 μM IP_6_ was added to the eluate and incubated for an hour to cleave the CPD and decahistidine tag before applying onto a 5 ml StrepTactin column (IBA) pre-equilibrated with buffer B. The column was washed with 30 CV of buffer B and eluted using buffer B supplemented with 5 mM desthiobiotin. Purified MprF was concentrated using a 100 kDa cut-off Amicon concentrator and injected onto a S200 Increase size exclusion column with buffer B as the mobile phase. The peak fraction was concentrated to 5mg/ml and used for further experiments. The steps in the purification of PaMprF with HRV3C construct was essentially the same as above, with only a decahistidine tag instead of the CPD domain and incubating IMAC eluate with HRV3C protease instead of IP_6_.

### MSP1E3D1 purification

The membrane scaffold protein MSP1E3D1 was purified according to the previous published protocol (Ritchie et al, 2009). Briefly, the recombinant plasmid MSP1E3D1 in pET28a was used to transform *E. coli* BL21 DE3 cells and transformants were grown in Terrific Broth medium supplemented with 50 μg/ml kanamycin and 2 mM MgSO_4_ at 37°C. Cells were induced at an OD_600_ of 1.8 with 1 mM IPTG and harvested after 3 hours using centrifugation. 20 g cells from 6 L culture were resuspended in 20 mM phosphate buffer pH 7.4 containing 1 % Triton X-100, 1 mM PMSF and 4 μg/ml DNase. Cell lysis was performed using sonication and debris were removed using centrifugation at 40,000 g for 40 minutes. The supernatant was applied to a 20 ml HisPrep column pre-equilibrated with 40 mM phosphate buffer pH 7.4. Three washes of 10 CV each were performed using the following buffer – (a) 40 mM Tris pH 8, 300 mM NaCl, 1% TritonX-100, (b) 40 mM Tris pH 8, 300 mM NaCl, 20 mM imidazole, 50 mM sodium cholate, (c) 40 mM Tris pH 8, 300 mM NaCl, 50 mM imidazole.

The elution was finally performed using 40 mM Tris pH 8, 300 mM NaCl, 200 mM imidazole. Recombinant TEV protease containing polyhistidine tag was added to the eluate and dialysed overnight against 25 mM Tris pH 7.4, 100 mM NaCl, 0.5 mM EDTA. The cleaved and dialysed protein was further purified through reverse IMAC and concentrated to 10 mg/ml and stored at −80°C till further use.

### Preparation of lipid stock

*E. coli* polar lipids (Avanti) in chloroform were dried in a glass test tube under a constant stream of argon. Traces of chloroform were removed using overnight desiccation. The dried lipids were resuspended in 50 mM Tris, 200 mM NaCl and 2 % CHAPS at a concentration of 20 mg/ml and sonicated at 25 kHz with 2 seconds on and 6 seconds off cycle using a microprobe, till the solution became clear. This took an approximate time of 5 minutes.

### Reconstitution of MprF in nanodisc

PaMprF was purified as above (after Ni-NTA affinity chromatography) and concentrated to 4 mg/ml. MSP1E3D1 and *E. coli* polar lipids in 2 % CHAPS were mixed at a ratio of 1:30 and incubated at RT for 30 minutes before placing on ice. MprF was added to the mix such that the ratio of MprF, MSP1E3D1 and lipids was 1:4:120 and incubated on ice for 15 minutes. An equal volume of Biobeads SM2 (BioRad) pre-equilibrated in Buffer A was added to the mix and allowed to stir gently overnight along with IP_6_ for removal of CPD tag. After detergent removal, the nanodisc mix was collected and centrifuged at 20,000 g to remove any precipitate before diluting 15 times. This was further purified with StrepTactin column. The eluate was then subjected to size exclusion chromatography on a Superose 6 Increase column. The peak fraction was concentrated to 2.5 mg/ml and used for further experiments.

### Cryo-EM grid preparation and Data Acquisition

The top side of Quantifoil Au R1.2/1.3 or UltraAufoil 1.2/1.3 (Russo & Passmore, 2014) (GDN) and Au R 0.6/1.0 (nanodisc) 300 mesh holey carbon grids were glow discharged at 25 mA for 60 seconds in air using a PELCO glow discharger (Ted Pella). 3 μl sample was applied on the grid and allowed to stand for 10 seconds at 16°C and 100% humidity, followed by blotting for 3-3.5 seconds with a blot force of 0 and subsequent vitrification in ethane using Vitrobot Mark IV (ThermoFisher Scientific). Data collection was performed with a 300kV Titan Krios G3i (ThermoFisher Scientific) with a Falcon 3 detector in counting mode at a nominal magnification of 75,000 x using the EPU software (ThermoFisher Scientific). The resulting pixel size was 1.07 Å and the image acquisition was performed at a defocus range of −1.5 to −3.0 μm. The dose was measured to be 0.55 e^-^/pixel/sec with a total exposure of 60 seconds and movies were fractionated into 25 frames resulting in ∼ 1.15 e^-^/ Å^2^/frame.

### Image Processing and Modelling and model refinement

The initial data processing steps were performed in Relion 4.0 (Kimanius et al, 2021; Zivanov et al, 2020; Scheres, 2012). Motion-correction was performed using Relion, followed by estimation of contrast transfer function with Gctf (Zhang, 2016). Autopicking was performed using Topaz (Bepler et al, 2019). For the data set in GDN, total of 1,239,668 particles were extracted with a box size of 320 pixels, and iterative 2D classification was used to clean up the data, yielding 597,348 particles. Initial model was generated and further 3D refinement with C2 symmetry was performed to obtain a map at a resolution of 4.17 Å. Multiple 3D classification steps were performed to obtain homogeneous particles. In the GDN dataset, classes with well resolved transmembrane helices were chosen for Bayesian polishing with 375,489 particles. These particles were then imported to cryoSPARC (Punjani et al, 2017) for further processing using non-uniform refinement and local and global CTF refinement to yield a map of ∼3.2 Å with C1 symmetry applied. Ab-initio reconstruction and heterogenous refinement was used to separate classes having only 1 or 2 soluble domains comprised of 168,060 and 109,157 particles respectively. The classes with both soluble domains intact were refined using non-uniform refinement with C1 or C2 symmetry. The final map sharpening was done with Relion PostProcess with automatic B-factor sharpening or with adhoc B-factor. Asymmetric refinement and reconstruction of classes containing a single soluble domain was also performed in cryoSPARC similarly to yield a map at 3.3 Å resolution.

The nanodisc dataset was collected at the same settings as above (1.07 Å sampling and Falcon 3 in counting mode) and had 3388 movie stacks that resulted in 1,701,373 particles. After multiple rounds of 2D classification, an initial model was generated using 428,167 particles. 3D classification was used to further remove heterogeneous classes and 138,067 particles were subjected to 3D refinement to yield a map at a resolution of 4.4 Å. After Bayesian polishing, the particles were exported to cryoSPARC for further processing. Non-uniform refinement in C2 symmetry yielded a map at 3.5 Å, which was sharpened in Relion with automated as well as adhoc B-factor sharpening. A subset of particles chosen from ab-initio and heterogenous refinement with no symmetry imposed resulted in a map of 3.6 Å resolution after non-uniform refinement.

The model for PaMprF was built in coot using maps that were sharpened with Relion (auto or adhoc B factor) or deepEMhancer or EMReady (He et al, 2023; Sanchez-Garcia et al, 2021). The crystal structure of soluble domain PDB – 4v35 was docked into the map using chimera and then built in coot (Emsley et al, 2010; Pettersen et al, 2004). The TM helices were modelled as polyalanine and the assigned sequence were verified with the predicted AlphaFold model. The lipid densities were identified by calculating FSC weighted difference (fofc) maps using servalcat (Yamashita et al, 2021). Refmac5/servalcat was used to generate the other monomer based on C2 symmetry. Model refinement was performed in real space refinement with Phenix or in reciprocal space with Refmac (Afonine et al, 2018; Adams et al, 2012; Murshudov et al, 2011; Murshudov, 2016). The figures were made with PyMOL, Chimera or ChimeraX (Pettersen et al, 2004, 2021; DeLano, 2002). The cavity analysis was performed using KVFinder web server (Oliveira et al, 2014) and the topology of PaMprF was made with PDBsum (Laskowski et al, 2018).

### Cysteine crosslinking

The wild type and double mutant membranes were thawed and diluted to a concentration of 1mg/ml using buffer A without β-mercaptoethanol. The membranes were treated with either DMSO (control) or 2 mM BMOE (bismaleimidoethane) (ThermoFisher-Pierce^TM^ Cat# 22323) and incubated for thirty minutes at room temperature. Samples were denatured using non-reducing SDS loading dye (samples containing 20 mM DTT were loaded separately) and SDS-PAGE gels were run, followed by transfer to a PVDF membrane. The membrane was probed using HRP conjugated antibody against Strep Tag and developed using ECL substrate.

## Acknowledgements

We acknowledge the National Cryo-EM facility, Bangalore, for data collection, which was supported by the Department of Biotechnology, DBT/PR12422/MED/31/287/2014, and the computing facility in the Bangalore Life Science Cluster in particular Mr Chakrapani. We thank all the lab members for their comments on the manuscript. SJ acknowledges the graduate fellowship from TIFR/NCBS. K.R.V. acknowledges the support of the Department of Atomic Energy, Government of India, Project Identification No. RTI 4006. KRV is part of the EMBO Global Investigator Network.

## Conflict of interest

The authors declare no conflict of interest.

## Data Availability

The cryo-EM maps and the coordinates have been deposited in the EMDB and PDB respectively with the following accession codes.

PaMprF in GDN, C2 symmetry – EMD-61238 and PDB 9J8Q

PaMprF in GDN, C1 symmetry – EMD-61239 and PDB 9J8R

PaMprF in nanodisc – EMD-61240 and PDB 9J8S

## Notes

### Competing Interest Statement

The authors have declared no competing interest.

