## Supplementary Information for "Structures of Multiple Peptide Resistance Factor from *Pseudomonas aeruginosa*"

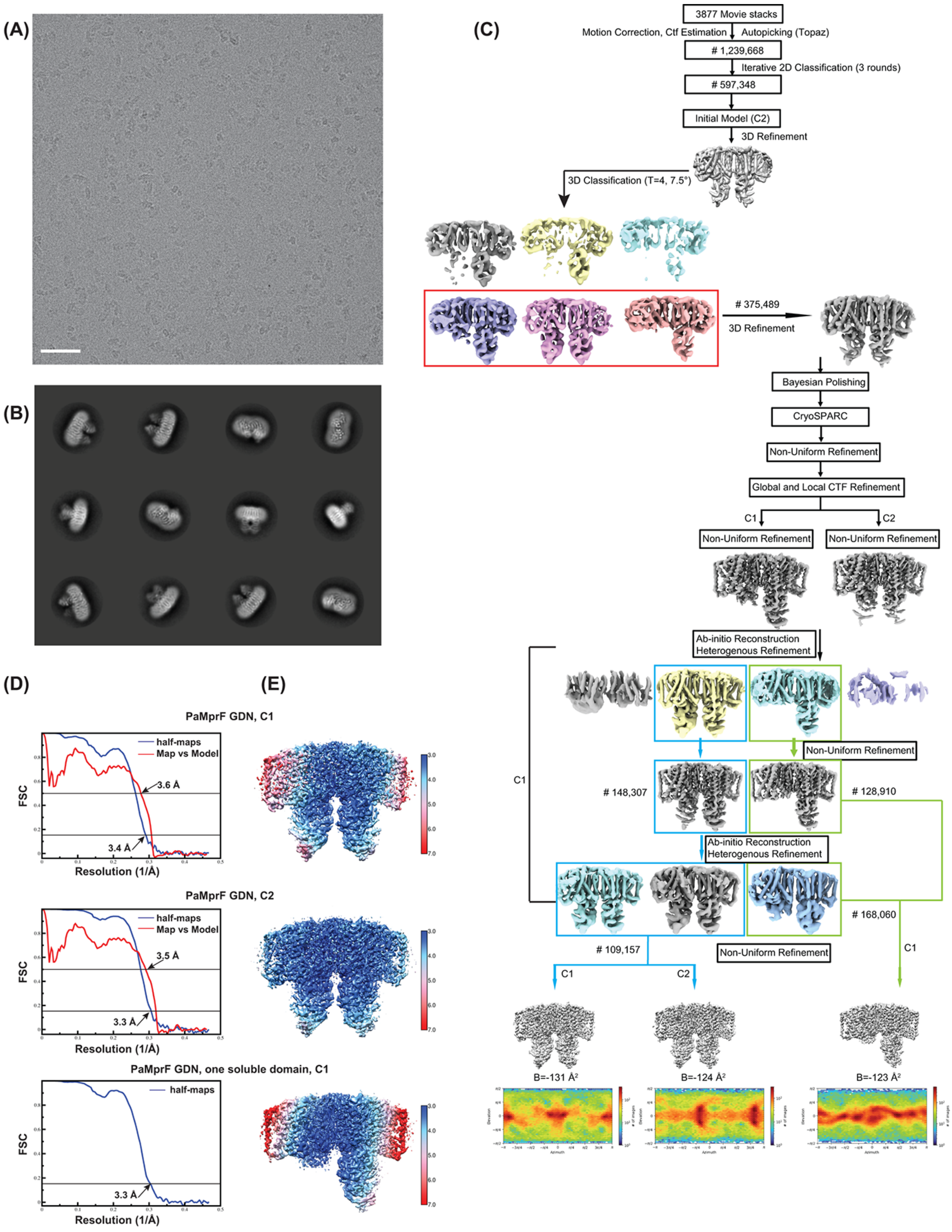


**Figure S1. CryoEM analysis of PaMprF purified in detergent** (A) Representative micrograph showing MprF particles. Scale bar is 500 Å. (B) Representative 2D class averages with a box size of 320 pixels or 342.4Å. (C) CryoEM data processing workflow for PaMprF in GDN. Both relion and cryosparc were used for image processing. The final sharpened maps and the orientation distribution plot of the PaMprF obtained from Cryosparc represented as a heatmap are shown at the bottom. (D) Fourier Shell Correlation curves of the cryoEM maps of PaMprF (GDN) with C1 and C2 refinement of the particles with both soluble domains ordered and the particles with only one soluble domain. The resolution is estimated from the FSCs by comparing the half-maps at 0.143 (blue colour) and the map vs model at 0.5 (red colour). Model docking and refinement was not performed for the map obtained with only one ordered soluble domain. (E) Local Resolution of PaMprF cryoEM maps as estimated by Relion. Colour key is given on the right with blue and red representing 3 Å and 7 Å respectively.

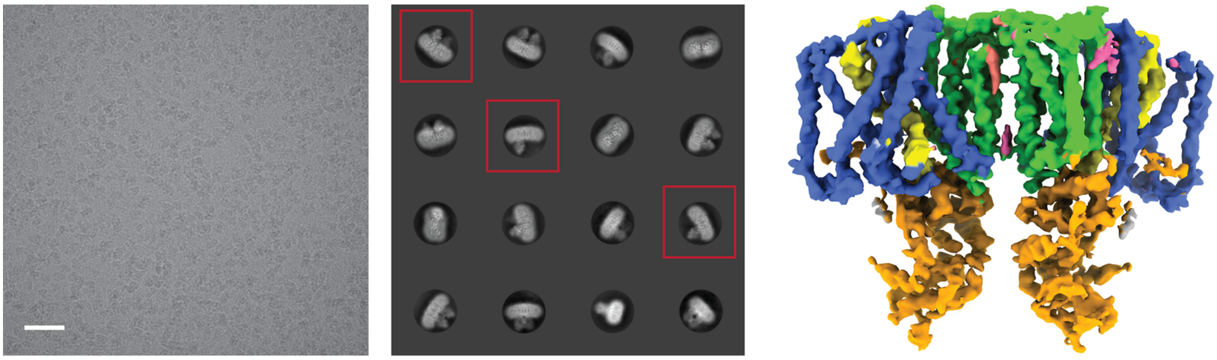

(A) (B) (C)

**Figure S2. CryoEM map of PaMprF expressed and purified with PreScission cleavage site and a poly-histidine tag.** (A and B), Representative micrograph and reference-free 2D class averages of PaMprF with a HRV 3C cleavage site. The scale bar in the micrograph is 500 Å and the box size of the 2D classes are 320 pixels (sampled at 1.07 Å/pixel). Few classes marked in red square show the populations with single soluble domain ordered and a common feature in both the constructs of PaMprF (with CPD and without CPD). (C) CryoEM map of PaMprF purified in GDN using a PreScission protease site and the tag was cleaved before the sample prep for cryoEM. The resolution of the map is ~ 4Å with C2 symmetry applied and coloured as in figure 1.

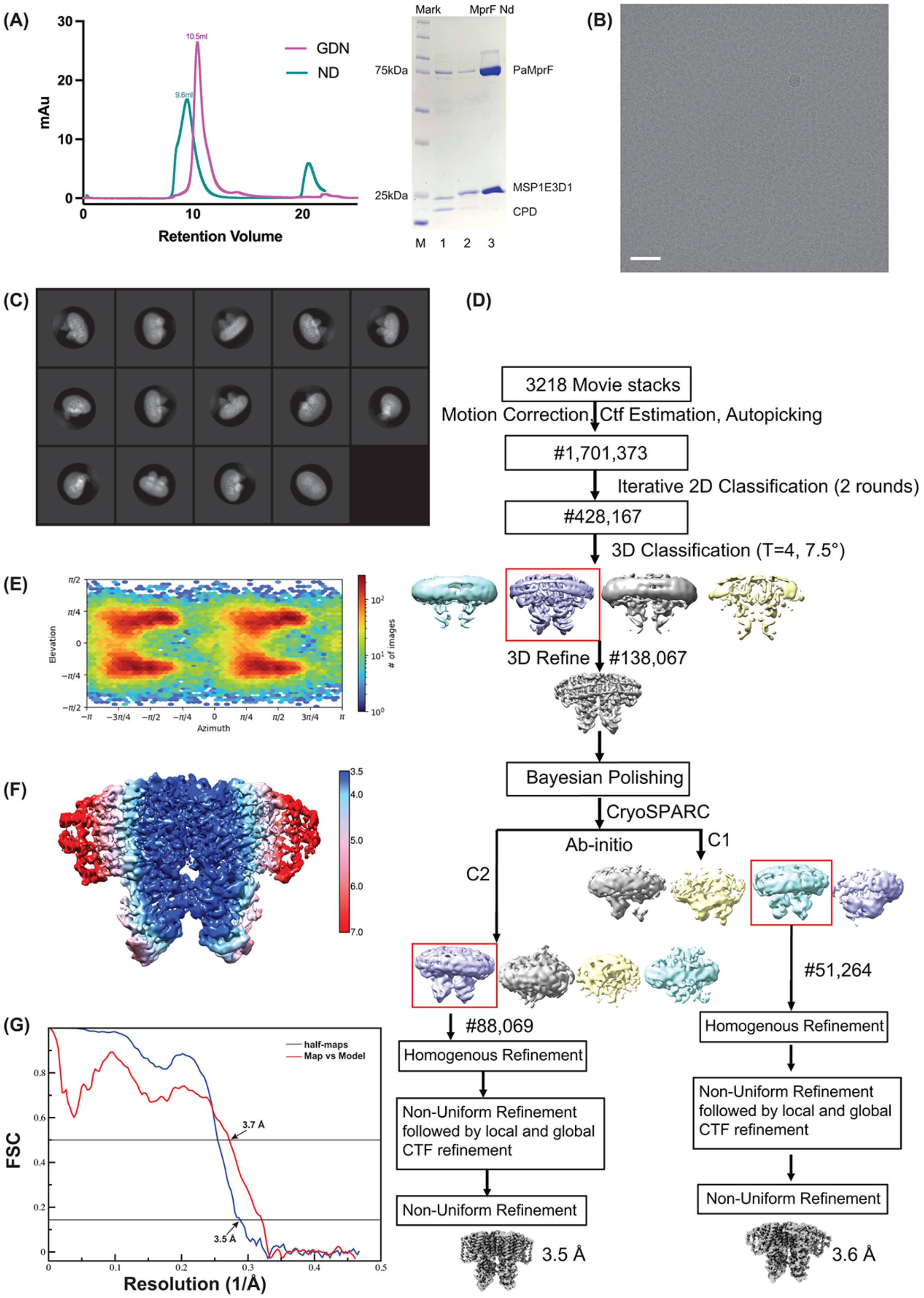


**Figure S3. CryoEM analysis of PaMprF purified in nanodisc** (A) Reconstitution of MprF in nanodisc. The size exclusion chromatography profile and the gel showing the enzyme and nanodisc protein indicative of successful reconstitution. When compared to MprF in GDN, the nanodsic is profile is slightly shifted and broad. A SDS PAGE gel showing the reconstitution of PaMprF in nanodisc. Lane M is marker, Lane 1 is PaMprF after Ni-NTA and reconstituted with MSP1E3D1 and addition of IP6 to remove the CPD tag, Lane 2- eluate from Strep affinity column to remove the CPD and his-tag and lane 3 – is the fraction after gel filtration and concentration showing both PaMprF and MSP1E3D1. (B) Representative micrograph showing the particles. Scale bar is 500 Å. (C) Representative 2D class averages of MprF in nanodisc with a box size of 320 pixels or 342.4 Å. (D) Image processing workflow of MprF in nanodisc. (E) Orientation plot of the PaMprF obtained from CryoSPARC represented as a heatmap (F) Local Resolution of PaMprF as estimated by Relion. Colour key shows the core of the enzyme is around 3.5 Å with the periphery and the nanodisc belt at lower resolution. (G) Fourier Shell Correlation curve of PaMprF (nanodisc). The resolution is estimated from the FSCs by comparing the half-maps at 0.143 (blue colour) and the map vs model at 0.5 (red colour) reveals 3.5 and 3.7 Å respectively.

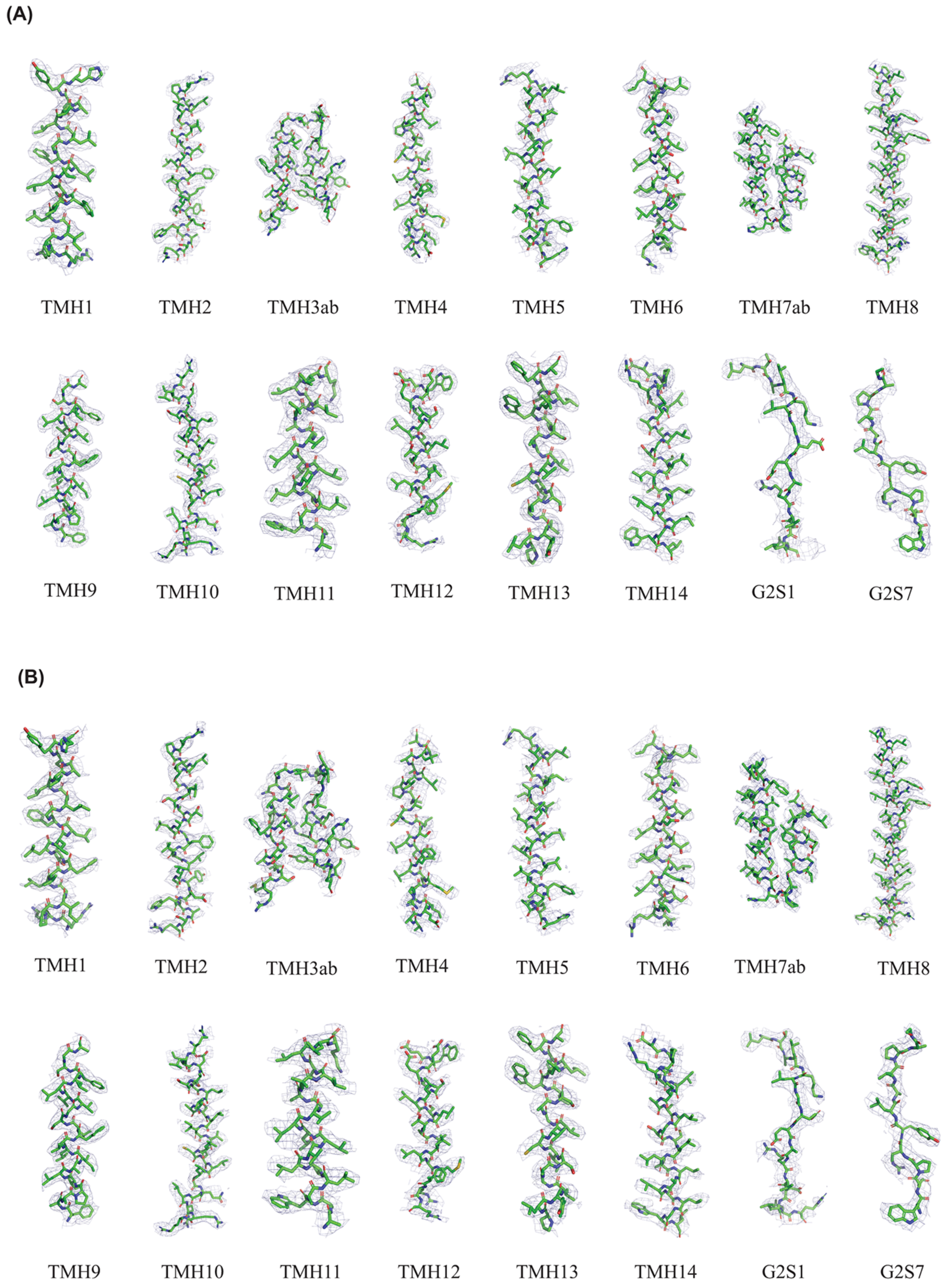


**Figure S4. Representative cryoEM density from the GDN and the nanodisc reconstructions.** Fit of the model to the cryoEM density of the various TM helices and β-sheets into the GDN (A) and nanodisc (B) maps. The model is shown in stick representation and the carbon atoms in green colour is encased in a blue mesh of the final sharpened cryoEM map contoured at 4 σ.

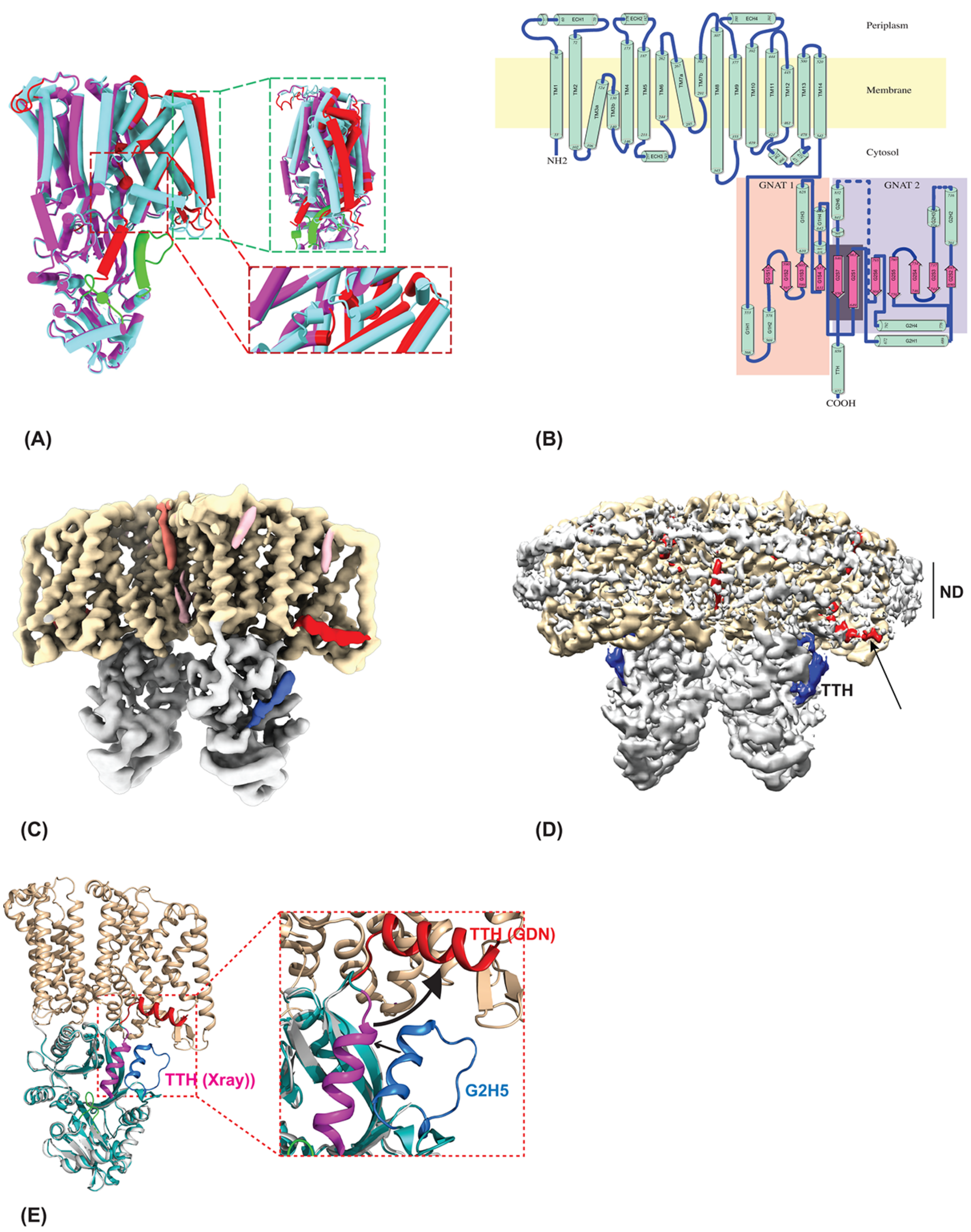


**Figure S5: Structural description of PaMprF.** (A) Comparison between experimental and AlphaFold predicted model of PaMprF. The predicted model by AlphaFold is shown in magenta and the experimental model in cyan. Region of the predicted model for which the density is ambiguous or not observed in the cryoEM map is coloured green. Regions of the predicted model where the rmsd between the Cα of the two models is > 3Å is coloured in red. The inset shows the side view of transmembrane helix 5 at the periphery (top) and the loop between the two re-entrant helices (bottom).

(B) **Topology diagram of PaMprF**. The TM helices are at the N terminus of the polypeptide chain. The secondary structural elements – α-helices and β-sheets are coloured in green and magenta respectively. The secondary structures are scaled to the length and the first and terminal residues are numbered. The membrane region is shaded in yellow and the TM helices are numbered as TMH1-14. The cytosolic domain constitutes two GNAT domains 1 and 2 (shaded in colour of salmon and purple). The helices and sheets of the cytosolic domain are named corresponding to the GNAT domain 1 or 2 from N to C-termini. For example, G2S1 corresponds to the second GNAT fold and first β-strand. The strands 5 and 11 are shared by both the GNAT domains. The topology diagram was generated by PDBsum server (Laskowski et al, 2018).

(C) **Unmodelled tubular density in the cryoEM map of PaMprF GDN.** An unsharpened map of PaMprF (GDN) containing both the soluble domains reconstructed using C2 symmetry. The TM domain, soluble domain and the TTH are coloured wheat, grey and red. The additional unmodelled density in the soluble domain is coloured in blue.

(D) **Unmodelled density in the cryoEM map of PaMprF nanodisc**. The cryoEM map of PaMprF in nanodisc sharpened with a B factor of -65 $Å^2$ is shown here with the TM and soluble domain coloured in wheat and grey. The density for TTH is shown in blue and the arrow marking the red density, which is currently unmodelled occupies a similar place as the TTH in GDN shown in panel C is most likely a lipid molecule. The nanodisc belt (grey) in the cryoEM map can be clearly seen in this map.

(E) **Comparison of models from full-length PaMprF (GDN) and crystal structure of transferase domain**. An overlay of monomer model of PaMprF (GDN) and the soluble domain of the X-ray structure of PaMprF (4v35). The region missing in the cryoEM PaMprF map and the model belonging to the helix 5 of second GNAT fold (G2H5) is coloured blue. The TTH in the soluble domain is coloured magenta, which is similar to the position of TTH in the nanodisc model. The inset on the right indicates the possible movement of G2H5 towards the membrane as shown here in the GDN model (helix in red).
